# Dynamic Disulfide Bond Topologies in von-Willebrand-Factor’s C4-Domain Undermine Platelet Binding

**DOI:** 10.1101/2022.08.20.504523

**Authors:** Fabian Kutzki, Diego Butera, Angelina J. Lay, Denis Maag, Joyce Chiu, Heng-Giap Woon, Tomáš Kubař, Marcus Elstner, Camilo Aponte-Santamaría, Philip J. Hogg, Frauke Gräter

**Affiliations:** Heidelberg Institute for Theoretical Studies, Schloß-Wolfsbrunnenweg 35, Heidelberg, Germany; University of Heidelberg, Im Neuenheimer Feld 205, Heidelberg, Germany; Institute of Physical Chemistry, Karlsruhe Institute of Technology (KIT), 76131 Karlsruhe, Germany; The Centenary Institute, University of Sydney, Camperdown, NSW, 2050, Australia

## Abstract

**Background:** The von Willebrand Factor (vWF) is a key player in regulating hemostasis through adhesion of platelets to sites of vascular injury. It is a large multi-domain mechano-sensitive protein stabilized by a net of disulfide bridges. Binding to platelet integrin is achieved by the vWF-C4 domain which exhibits a fixed fold, even under conditions of severe mechanical stress, but only if critical internal disulfide bonds are closed.

**Objective:** To quantitatively determine C4’s disulfide topologies and their implication in vWF’s platelet-binding function via integrin.

**Methods:** We employed a combination of classical Molecular Dynamics and quantum mechanical simulations, mass spectrometry, site-directed mutagenesis, and platelet binding assays.

**Results:** We quantitatively show that two disulfide bonds in the vWF-C4 domain, namely the two major force-bearing ones, are partially reduced in human blood. Reduction leads to pronounced conformational changes within C4 that considerably affect the accessibility of the RGD-integrin binding motif, and thereby impair integrin-mediated platelet binding. Our combined approach also reveals that reduced species in the C4 domain undergo specific thiol/disulfide exchanges with the remaining disulfide bridges, in a process in which mechanical force may increase the proximity of specific reactant cysteines, further trapping C4 in a state of low integrin-binding propensity. We identify a multitude of redox states in all six vWF-C domains, suggesting disulfide bond reduction and swapping to be a general theme.

**Conclusion:** Overall, our data put forward a mechanism in which disulfide bonds dynamically swap cysteine partners and control the interaction of vWF with integrin and potentially other partners, thereby critically influencing its hemostatic function.

**Essentials:**

- Platelet integrins interact with the disulfide-bonded C4 domain of von Willebrand Factor
- The redox state of vWF-C4’s disulfide bonds is studied by molecular simulations and experiments
- Two bonds are reduced causing C4 unfolding and disulfide swapping
- Opening of disulfide bonds impairs integrin-mediated platelet binding

## Introduction

The von Willebrand Factor (vWF) is a large mega-Dalton multimeric extracellular protein essential in achieving and regulating primary hemostasis [1]. Its mature monomers consist of 2050 amino acids, divided into 12 heavily disulfide-bonded protein domains (Fig. 1A). It features 169 cysteines across its sequence [2], which are considered to be mainly paired in vWF multimers. Cysteine-derived disulfide bridges between the carboxy- and the amino-terminal domains establish vWF dimers and multimers, respectively [1]. Other disulfide bonds protect the structural integrity of individual domains [1].

**Figure 1:**
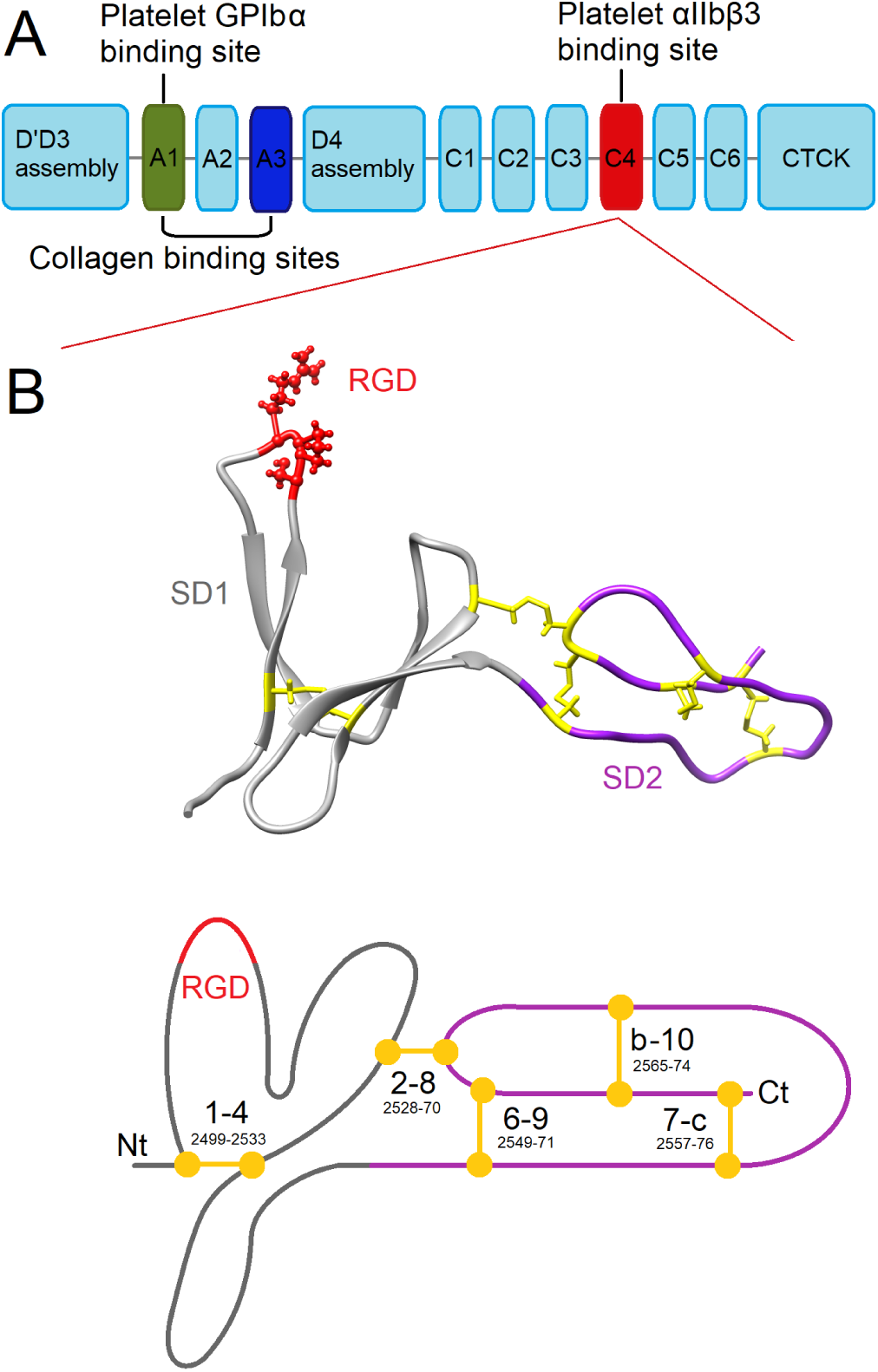
vWF-C4 domain: A: Scheme of the vWF monomer, highlighting the position of the C4 domain. Platelet GPIbα, integrin αIIbβ3, and collagen binding domains are indicated. B-up: Cartoon representation of vWF-C4 (85 residues, 2493-2577 of the vWF sequence) according to its NMR structure [21]. The C4 integrin-binding motif (RGD, residues 2507-2509) and the five disulfide bonds are represented as sticks and spheres, the two sub-domains (SD1 and SD2) are differentiated by color. B-down: Scheme of the vWF-C4 domain labeling the five disulfide bonds according to the overall vWF-C domain topology [57] (amino acid id.s are also indicated).

There are two essential functions vWF carries out in hemostasis, namely, the adhesion of platelets to collagen at sites of vascular injury as well as the transportation and the half life increase of Factor VIII through protective binding [1]. Mutations in the structure of vWF or dysregulation of its concentration in the blood stream lead to severe, and often life threatening bleeding disorders, a medical condition with a broad range of phenotypic responses ranging from acute hemorrhaging to thrombus formation, known as von Willebrand disease [3, 4]. Many of the known disease mutants are mutations of cysteines, highlighting their essential role in vWF function [5].

Triggered by the shear stress of the flowing blood, vWF initiates the primary response to injury by recruiting platelets to the collagen matrix of the endothelium [3, 6]. Under conditions of low hydrodynamic forces, vWF adopts a condensed globular structure that gets elongated upon surpassing shear rates at the scale of 10^3^ s^*-*1^ [6]. Here, the vWF A1 and A3 domains connect the vWF multimers to collagen [1] (Fig. 1A) and vWF A1 binds to platelets via the interaction with the platelet glycoprotein Ib*α* receptor [3]. The tensile forces acting on vWF multimers, in dependence of shear rate and vWF structure elongation, increase the binding affinity between A1 and GPIbα [7].

In contrast to these non-covalent force responses of vWF, the dynamics of the covalent binding states given by the disulfide connectivity, as a response to tensile forces from flowing blood, remain largely unknown. It is established that disulfide bonds drive vWF multimerisation and protect it from unfolding under the influence of shear in the flowing blood [1]. Consequently, improper disulfide pairing is related to dimerization-[8, 9] and multimerization defects [10], as well as to the structural rearrangement and gain of function of the domain A1 [11]. Beyond conferring structural integrity, disulfide bonds have been recently discovered to directly participate in hemostatic functions. Thiol disulfide shuffling has been suggested to mediate vWF multimer size [12], oligomerization [13, 14], and platelet binding [15]. Moreover, auto-inhibition of vWF for the binding of platelets, initiated by A1-A2 inter-domain interactions, is controlled by a disulfide bond switch in the vWF A2 domain [16]. In addition, blockage of free thiols in vWF has been reported to interfere with its binding to collagen [17]. Beyond vWF, other extracellular proteins involved in hemostasis have been shown to be mechano-redox controlled. The shear-dependent (de)adhesion of integrin αIIbβ3 to fibrinogen is an example of this [18]. Fibrinogen by itself exists in a multitude of force-responsive covalent forms which dynamically exchange to drive fibrin polymerisation [19].

In case of vWF, while the importance of cysteine redox states and disulfide connections has been recognized, structural data on the major disulfide bonded domains C and D have only very recently become available [14, 20–22]. Hence, a mechanistic and quantitative view on redox regulation and the role of force therein is currently lacking. We here focus on resolving the atomistic principles of disulfide bond reduction of the vWF-C4 domain. C4 plays a direct adhesive role as an anchoring point for platelets [23] and is the only C-domain, along with CK [24], whose structure has been resolved [21]. C4 is a small 85-amino acid domain composed of two flexible sub-domains SD1 and SD2 and is structurally stabilized by five disulfide bridges (Fig. 1B). It contains an integrin-binding RGD motif at the tip of the first beta-hairpin in the SD1 subdomain where it can connect to the platelet integrin receptor αIIbβ3 [3]. The C4 domain and most importantly the loop containing the RGD motif are topologically protected from mechanical unfolding by a set of disulfide bonds [21].

In this study, we asked if the redox states of inter-cysteine disulfide bonds in C4 dynamically regulate the presentation of RGD to platelets, which is one of the key functions of vWF. By integrating molecular dynamics (MD) simulations and quantum mechanics/molecular mechanics (QM/MM) calculations with mass spectrometry and platelet binding assays, we quantitatively demonstrate the existence of two reduced states in C4 in human blood. We find these reduced states to impair the adhesion of platelets to vWF by compromising the accessibility of C4’s RGD motif to integrin. The free thiols propel disulfide bond exchange, resulting in swapped linkages, again with functional implications. Moreover, as molecular principle guiding this process, we find that force could be beneficial to produce these reduced states and to enable subsequent swapping. Overall, our study provides new evidence to the emerging concept of chemo-mechanical sensitivity in bio-molecules and the regulative effect this has on vWF’s hemostatic function.

## Methods

### Redox states of human vWF-C domain disulfides

To determine the redox state of human vWF-C domain disulfides, blood was collected by venesection from 10 healthy donors on no medications (5 male, 5 female, 18-57 years old). Liquid chromatography, mass spectrometry and data analysis were performed. Cysteine-containing vWF peptides were analysed (Fig. 7 and Table S1). The different redox forms of the cysteine residues were quantified from the relative ion abundance of peptides labelled with ^12^C-IPA and/or ^13^C-IPA (Fig. 5). The data was searched for peptides containing free cysteine thiols and these were not detected, which indicates that alkylation of unpaired cysteine residues by ^12^C-IPA or ^13^C-IPA was complete in the proteins.

### Cloning and expression of human vWF

Cysteine to Alanine substitutions within the vWF-C4 domain at positions 2499 and 2533, and 2528 and 2570 were generated in full-length human vWF DNA using a Site-directed Ligase Independent Mutagenesis (SLIM) approach [25]. The C2528A and C2570A mutants were created sequentially with C2528A being created first from a wild-type vWF template, followed by the fabrication of the C2570A mutation using the C2528A mutant as a template. Both mutations were confirmed by DNA sequencing.

### Platelet adhesion studies

To investigate the consequences of these substitutions for vWF-C4’s ability to bind platelet integrin, a platelet adhesion stduy was performed. Washed human platelets were perfused over vWF matrix using polydimethylsiloxane (PDMS) microfluidic channel devices. Platelets were perfused over the vWF-coated matrix at a shear rate of 1000 s^*-*1^. Platelet adhesion was recorded microcopically. Images were captured at 1 s/frame for 3 min. Platelets adhering to the matrix for at least 2 s were considered adherent platelets. Rolling platelets were defined by their movement of >1 cell diameter in 10 s following tethering to the matrix. Platelets moving <1 cell diameter in 10 s were considered stationary platelets. The kinetics of platelet adhesion was analysed at 30 s intervals.

### Mapping of disulfide bonds in human vWF-C domain

To map disulfide bonds in human vWF-C domains, Peptides were ionized by electrospray ionization at +2.0 kV and analysed on a Q-Exactive Plus mass spectrometer using collision-induced dissociation fragmentation. Disulfide-linked peptides were searched against human vWF sequence using Byonic analysis software with a false discovery rate of 1 %. Disulfide-linked peptides were manually inspected for accuracy. Precursor mass tolerance and fragment tolerance were set at 10 ppm and 0.6 Da, respectively. Variable modifications were defined as oxidized Met, oxidized cysteine (mono, di and tri) and glutathionylated cysteine with full trypsin and chymotrypsin cleavage of up to three missed cleavages.

### Molecular Dynamics Simulations

Different conformations of the C4 domain (fragment 2493-2577 of the vWF sequence, i.e. 85 residues) were taken from previous MD simulations [21] in a randomised fashion. We removed individual inter-cysteine disulfide bonds, resulting in 6 different cases: one with the protein fully oxidised (fo) and five more with one of the disulfide bonds removed (1-4, 2-8, 6-9, 7-c, b-10) (Fig. 1, 2). At least ten different starting conformations were considered for the fo as well as for any of the reduced cases. MD simulations were carried out starting from each one of these replicas for simulation lengths ranging from 140–210 ns for a cumulative simulation time of at least 1400 ns per system. Subsequently, we extracted randomised frames from these equilibrium simulations to conduct force-probe MD simulations under the influence of external pulling forces (50 pN, 100 pN, and 500 pN), applied to the termini of the protein (see Fig. 2). N=10 pulling simulations were carried out for each force and disulfide state, with variable duration from 70–200 ns. In addition, for the 2-8 system, simulations at the low forces of 50 pN and 100 pN were started from fully stretched configurations extracted from the 500 pN simulation (n=10 replicas of at least 100 ns for each force value). For each force and oxidation state a cumulative simulation time of at least 1000 ns was allocated.

**Figure 2:**
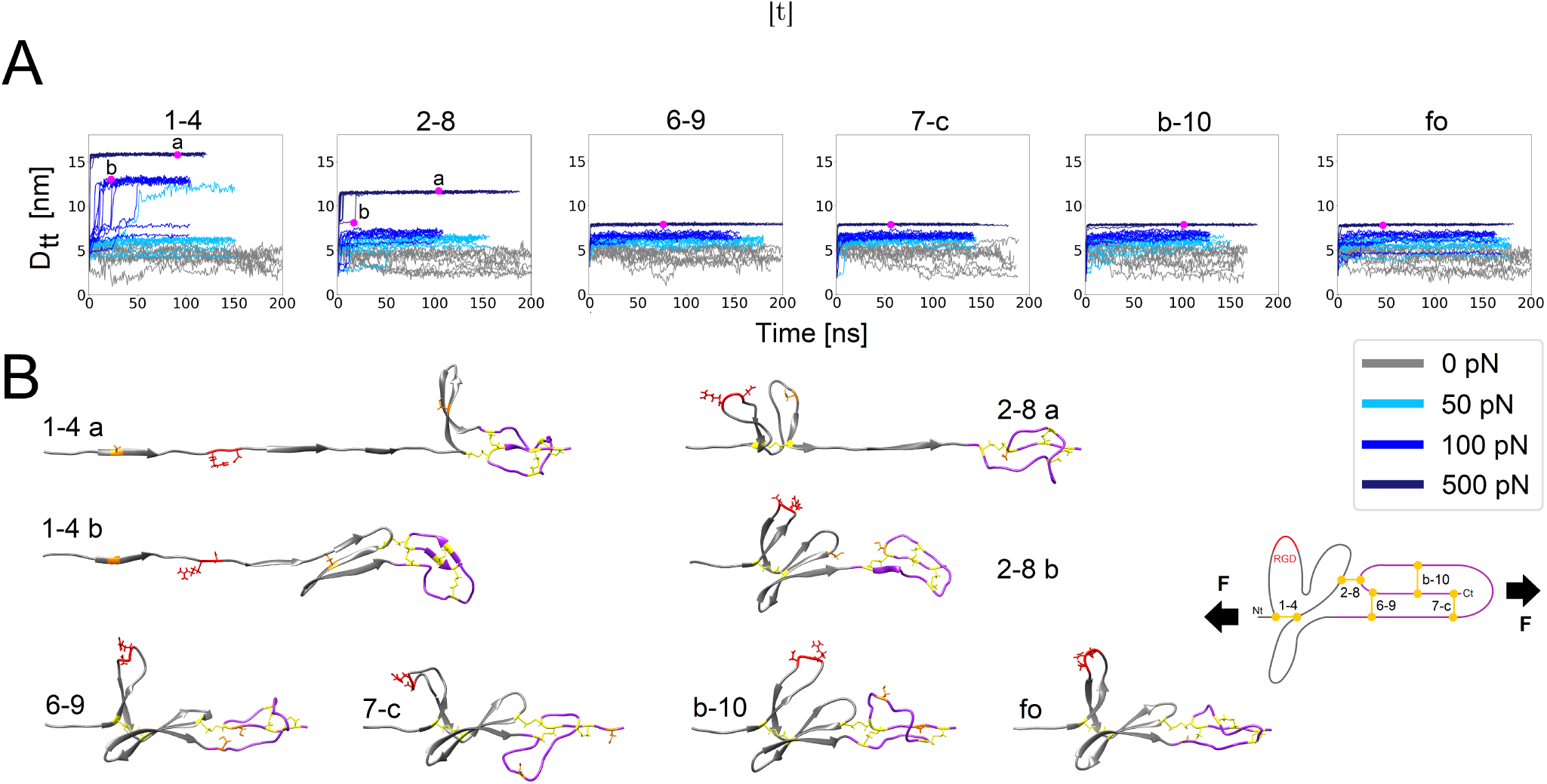
Tension-induced elongation of the vWF-C4 domain upon disulfide bond reduction. A: Elongation of the C4 domain under force was measured by the distance between termini (D_tt_), in the fully oxidised case (fo) and after reducing the indicated disulfide bond upon application of a constant force F (different force values depicted with different color). B: Examples of structures at the instants indicated with the magenta dots in A are displayed (SD1 region: gray; SD2 region: purple; RGD motif: red; disulfide bonds: yellow). At the right a scheme of the protein highlighting the disulfide bonds is included.

### QM/MM calculations

We included QM/MM calculations using semi-empirical Density-Functional Tight-Binding (DFTB) as QM method. Seven structures with high mutual RMSD values (larger than 0.43 nm) were selected from the Force-probe MD runs to serve as initial structures for QM/MM metadynamics simulations in which, after initial equilibration, the sidechains of CYS 1, CYS 2 and CYS 4 were made able to interact with each other, allowing them to undergo chemical reactions. Termini were subjected to a pulling load of 166 pN and the three sulfur distances were employed as general reaction coordinates for the metadynamics scheme. Gaussian bias potentials were added every 500 fs. Each simulation involved 16 walkers simulated for at least 18.75 ns at 300 K. The initial Gaussian height was set to 1.0 kJ/mol, the width to 0.2 Å and the bias factor to 50.

See experimental and simulation details in the supporting information, which includes references 25–56.

## Results

### Tension-dependent redox states of the vWF-C4 domain

C4 contains five disulfide bonds, which have been observed closed in NMR structures of this domain [21] (Fig. 1B). Fully oxidised, these bonds structurally stabilise C4, ensuring its ability to withstand the external shear force of the blood stream and thereby guaranteeing its capacity to optimally bind to platelets via the interaction with integrin. Reduction of these bonds may alter the structural integrity of the C4 domain affecting integrin (and thus platelet) binding. Disulfide bond reduction would leave free thiols as reaction products opening up the possibility of disulfide exchange.

Hence, we first investigated which effect the redox state of the disulfide bridges has on the conformation of the vWF-C4 domain under tension. We performed at least 10 force-probe MD simulations for each individual redox state (including the fully oxidised configuration, fo) by pulling the termini of C4 away from each other, exerting different forces of ∼50, ∼100 and ∼500 pN (Figure 2). We monitored the separation between the termini as a function of the protein elongation (Figure 2A). As expected, due to the disulfide bonds, the fully oxidised structure practically did not elongate upon force application, i.e. only the termini were stretched, resulting in a maximum inter-termini separation of ∼7.9 nm (for a pulling force of 500 pN). Opening the disulfide bonds which are located solely within the SD2 sub-domain, namely, 6-9, 7-c or b-10, resulted in similar elongation lengths as for the fully-oxidized protein. At lower forces and within the simulation time-scale of ∼100 ns, only intermediate elongations of ∼12.9 nm were observed upon reduction of the 1-4 bond (eight cases at 100 pN and one case at ∼50 pN out of n=10 simulations for each force). Here, application of force led to a complete flattening of the beta hairpin containing the RGD motif but it was not sufficient to completely open the other beta hairpin of SD1, namely the one carrying the bond 2-8 (state 1-4b in Figure 2). For 2-8, at a lower force regime only a residual stretching of the termini was observed, with an elongation similar to the fully oxidised case state 2-8b in Figure 2. The strength of the beta strand within SD1, formed in between sulfurs 4 and 6, was sufficient to protect the structure from unfolding even in the absence of bond 2-8. These simulations in the 50–100 pN regime already gave hints into the possible force-induced unfolding events for the reduced forms of C4, specially upon removal of 1-4. However, the simulation time-scale (∼100 ns) limited the observation of full force-induced opening conformational transitions. To overcome this issue, we applied a higher force of 500 pN. In this case, reduction of either the bond 1-4 in SD1 or the bond 2-8 which connects SD1 and SD2 triggered major unfolding events, resulting in final elongations of ∼15.9 nm and ∼11.5 nm for 1-4 and 2-8, respectively (comparable to those observed in previous simulations [17]). In this case, the SD1 sub-domain fully opened after reduction of 1-4 (state 1-4a in Figure 2) while it fully separated from the SD2 sub-domain upon removal of the 2-8 bond (state 2-8a in Figure 2). Thus, force impacts the conformation of reduced 1-4 and 2-8 C4 domains.

### C4 disulfide bonds 1-4 and 2-8 are partially reduced

The 1-4 and 2-8 bonds align themselves along the direction of the pulling axis, while 6-9, 7-c and b-10 adopt a more perpendicular orientation to this axis (see f.o. state in Figure 2B). Furthermore, reduction of these two bonds induced large unfolding of C4. Thus, we hypothesized that for vWF-C4 in human blood, 1-4 and 2-8 could partially exist in a reduced state. Remarkably, the cysteines bridged by these bonds have been suggested to also exist as free thiols before [15]. To further quantitatively test our hypothesis, we investigated their redox state in plasma of ten healthy human donors *ex vivo*, using mass spectrometry (Figure 3A). Although the two bonds were predominantly oxidized, a small fraction of the 1-4 and the 2-8 bonds were on average 3.4 and 2.7% reduced in the population of the vWF molecules. This result indicates that the C4 domain disulfide bonds 1-4 and 2-8 indeed have the potential to be labile. We were not able to detect reduced peptides for the other three disulfide bonds, but can not fully exclude their presence in human blood. Taken together, we have experimental evidence for two partially reduced disulfide bonds in C4, 1-4 and 2-8, which is consistent with [15].

**Figure 3:**
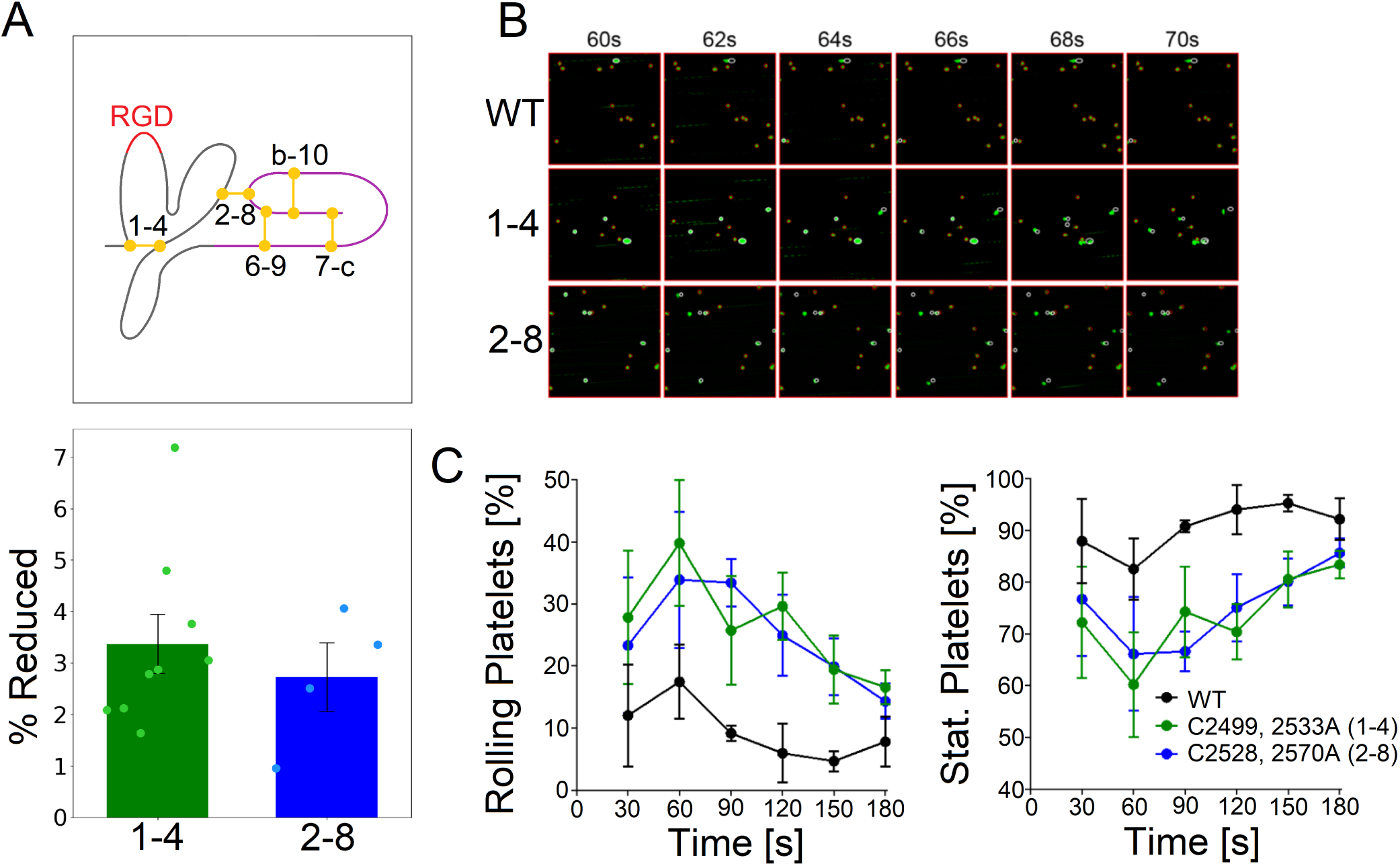
Disulfide bonds 1-4 and 2-8 in the vWF-C4 domain are partially reduced in blood and their genetic ablation impairs engagement of the C4 RGD motif with platelet integrin under fluid shear conditions. A: Reduction state of C4 disulfide bridges in healthy human donors. Shown is the percentage of reduced bonds for each patient (dots), the mean value of all patients (green and blue bars) as well as the standard deviation on the mean (black error bars). See above the scheme of the C4 indicating the location of these two bonds. B: Washed human platelets were perfused over vWF matrices at a shear rate of 1000 s^-1^ for 3 min. Examples of rolling and stationary platelets between 60 and 70 s of perfusion are shown for the wild-type (WT) protein and for the variants with either the 1-4 or the 2-8 removed by alanine substitution of the cysteines. C: Percentage of rolling and stationary platelets as a function of time of perfusion is displayed (mean ± S.E.M. of 4 biological replicates).

### Genetic ablation of either 1-4 or 2-8 bonds impairs platelet immobilization

In blood, the strained 1-4 and 2-8 bonds also exist in a reduced state (Fig. 3A) and C4 undergoes pronounced unfolding transitions under such conditions (Fig. 2). These two observations led us to ask if the C4 disulfide lability has functional consequences. To this end, we examined the effect of ablating these bonds on platelet adhesion. Accordingly, the 1-4 or 2-8 disulfides were eliminated by mutating both cysteines of the bond to alanines. The wild type and mutant proteins were expressed in HEK cells and collected from conditioned medium. The redox state of the C4 domain disulfides of the recombinant proteins was measured to ensure that elimination of either one of the C4 bonds did not change the redox state of the other one, or other C domain bonds in general. The disulfide status of recombinant wild-type and C4 domain disulfide mutants was very similar (Fig. S1). Moreover, the redox state of the recombinant proteins was comparable to human plasma vWF (compare Fig. 3A and S1). Thus, mutations did not alter the redox state of other C domains. Washed human platelets were perfused over vWF matrices at a shear rate of 1000 s^*-*1^ and the number of stationary and rolling platelets was measured at 30 s intervals (Fig. 3B). Elimination of either the 1-4 or 2-8 vWF-C4 disulfide bond resulted in significantly more rolling and fewer stationary platelets (Fig. 3C). Therefore, the 1-4 and 2-8 vWF-C4 disulfide bonds need to be intact for efficient engagement of the C4 RGD motif with platelet *α*IIb*β*3 integrin.

Our MD simulations provided a molecular explanation for this reduction in platelet binding affinity. Without the stabilising 1-4 bond, the beta hairpin containing the integrin binding RGD motif completely flattens (Fig. 4A). From a geometric standpoint, this conformation is not likely to be optimal for binding as indicated by the beta-hairpin opening-angle and Ramachandran plot for the three amino acids of the RGD motif. Application of an external force increased the probability of opening the RGD-containig beta hairpin (see non-zero probability for the angle *θ* ∼ 180^*o*^ when the force was not zero in Fig. 4A, left). In addition, we compared the RGD-motif backbone torsion angles observed in the simulations (after full expansion) with those measured in X-Ray structures of RGD-containing ligands bound to integrins. The angles observed in the X-Ray structures are close to those sampled in the force-free simulations, but this situation changes upon application of force. Especially the ARG and ASP angles calculated in the simulations increasingly distribute far from the experimental references as the pulling force increases. Considerable deformation of the RGD motif thus undermines optimal binding of integrin (and thereby of platelets) to the vWF mutant lacking the 1-4 bond.

**Figure 4:**
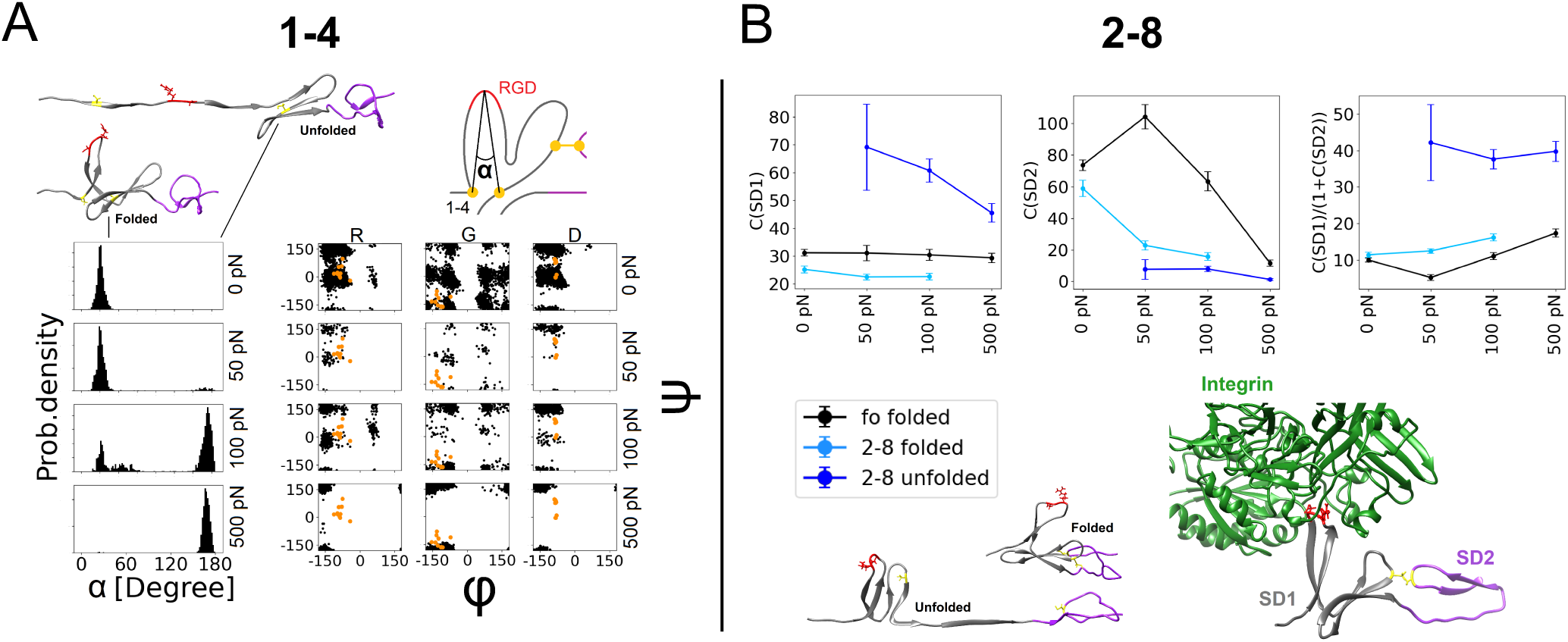
Alteration and accessibility of the RGD motif under force reduces binding to integrin of 1-4 and 2-8 reduced vWF-C4 variants. A: RGD beta hairpin angle α formed by Cys2499 (1), Gly2508 of the RGD domain, and Cys2533 (4) (see scheme) as well as Ramachandran angles ϕ and Ψ for the RGD motif (ARG-GLY-ASP) were recovered from simulations with the bond 1-4 reduced, under different forces (different horizontal panels). Cartoons at the top exemplify the folded (α ∼ 30°) and unfolded conformations (α ∼ 180°) of C4 (SD1: gray; SD2: purple; RDG motif: red, and 1 and 4 cysteines: yellow). 13 orange dots in the Ramachandran plots represent angles taken from the structures of RGD peptides in complex with integrin [58]. The Ramachandran angles for the force bearing cases were computed from simulations at the indicated forces but started from a fully elongated conformation of C4. B: A standard binding configuration between vWF-C4 (grey and purple) and αIIbβ3 integrin (green) was predicted by alphafold (bottom-right cartoon). Configurations of fully oxidised C4 as well as its variant with bond 2-8 opened were overlaid with this prediction, fitting the positions of the RGD beta hairpin. For the 2-8 reduced case, both states before and after full-unfolding were considered. For the low forces of 50 and 100 pN, these unfolded conformations were extracted from simulations that started from a pre-stretched conformation. Number of clashes, C, of the heavy atoms of C4 with those of Integrin were calculated separately for SD1 (left panel) and SD2 (middle panel). C is shown as a function of the applied force. A clash was assigned if the inter-atomic distance was below 0.3 nm. The clashes of sub-domain SD1 were then weighted by those of SD2, C(SD1)/(1 + C(SD2)), to thereby exclude conformations displaying large overlap between the SD2 domain and the integrin complex (right panel). In all cases, time-averages ± S.E.M. are presented.

The same argument cannot be applied to the mutant with bond 2-8 eliminated. Reducing this bond displayed no major effect on the conformation of the RGD binding beta hairpin (Fig. 4B). In this case, the existent 1-4 bond is sufficient to stabilise the RGD beta hairpin. However, as soon as the connection between the second beta hairpin of SD1 (carrier of cysteine 2) and the sub-domain SD2 is broken, an increased propensity to occlude the RGD binding beta hairpin is observed, according to our structural analysis as follows. We predicted the conformation of the complex formed by *α*IIb*β*3 integrin and the vWF-C4 domain using alphafold [42, 43] (Fig. 4B). Thereafter, we fitted all simulation frames of the fully oxidised case as well as the 2-8 reduced conformation to the predicted structure, by overlaying the RGD beta hairpin. Our original simulations for the 2-8 case, did not produce fully-expanded configurations for the 50–100 pN regime, thus we continued randomly chosen configurations from the 500 pN simulations at these lower forces. We measured clashes between backbone atoms of integrin and heavy atoms of the vWF-C4 sub-domains individually. In general, the unfolding of C4 as a result of the reduction of bond 2-8 increased the number of clashes between SD1 and the predicted position of integrin, while it reduced that number for SD2 (Fig. 4B, left and middle panels). The latter effect is especially pronounced for high pulling forces. We interpret the reduced number of clashes between SD2 and integrin as a geometrical effect that force has on the overall position of SD2, locating this sub-domain at a distant position relative to the integrin and preventing the overall sub-domain hinge motion of C4 as described in earlier works [21]. To test if the increased number of SD1 clashes upon unfolding is indeed the consequence of special SD1 conformations and not the result of single, particularly bad fits, we weighed the clashes of SD1 by the number of clashes of SD2 (plus one). Accordingly, structures presenting few SD2-integrin clashes (which generally indicate a reasonable fit) were preferred. This choice resulted in an even more pronounced effect, leaving the average number of SD1 clashes more than twice as high for the unfolded case, compared to both folded ones (Fig. 4B, right panel). From this analysis, we suggest 2-8 reduction hampers integrin–vWF C4 binding by steric hindrance.

### C4 undergoes disulfide bond shuffling

Reduction of 1-4 and 2-8 bonds leaves free thiols behind which may get in close proximity and thereby attack still-formed disulfide bonds. We tested this hypothesis *ex vivo*. vWF was immunoprecipitated from healthy donor plasma, digested with chymotrypsin and trypsin, peptides resolved by HPLC and analysed by mass spectrometry (see methods for details). The C4 domain ASPENPCLINECVR peptide containing the CYS2528-CYS2533 (2-4) disulfide bond was identified with very high confidence (p = 0.00066) (Fig 5). In addition to this new bond formed inside C4, the free cysteine 1 (C2499) established a bond with the cysteine C2494 (C3-10) of the neighbor C3 domain, as reflected by identification of the CLPSACEVVTGSPR peptide (Fig. S2). In consequence, disulfide bonds undergo intra-C4 and C3-C4 inter-domain disulfide bond swapping.

**Figure 5:**
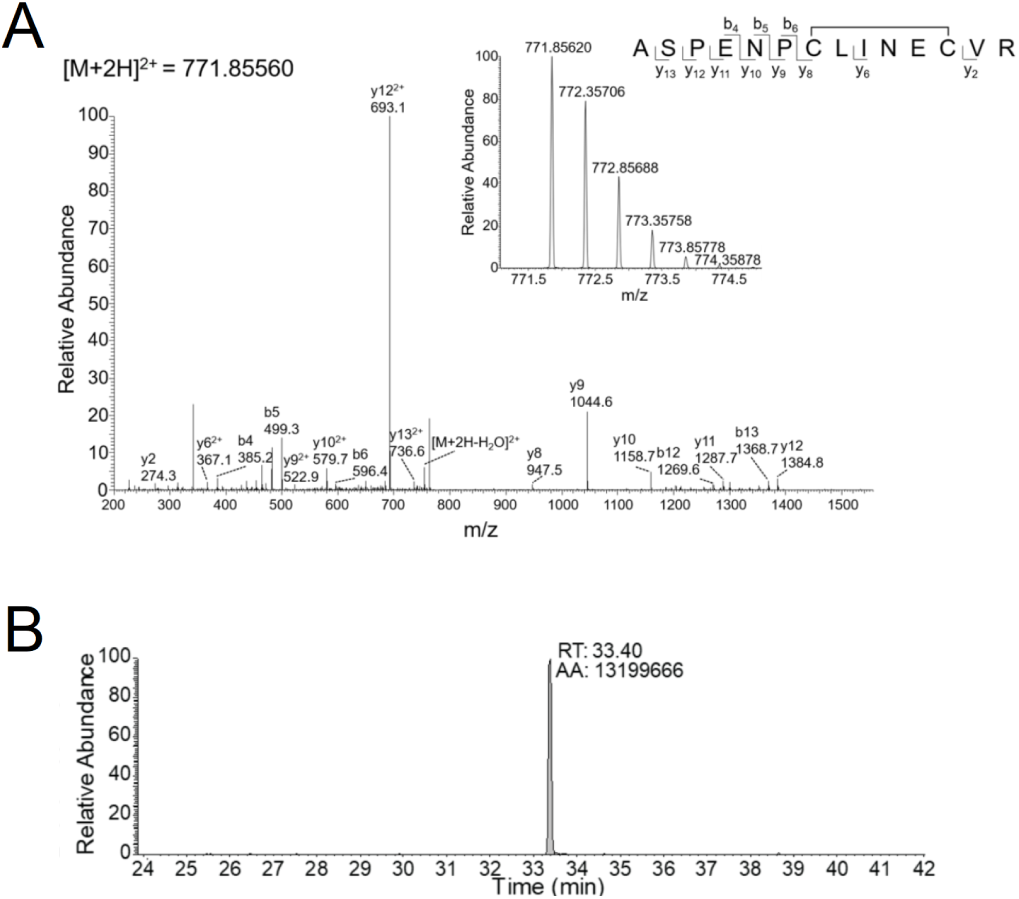
vWF-C4 undergoes disulfide bond shuffling: A: Representative tandem mass spectra of the ASPENPCLINECVR peptide. The accurate mass spectrum of the peptide is shown in the inset (observed [M + 2H]2+ = 771.85620 m/z and expected [M + 2H]2+ = 771.85560 m/z). B: HPLC resolution of the C4 domain ASPENP<u>C</u>LINE<u>C</u>VR peptide containing the C2528-C2533 (2-4) disulfide bond (cysteines underlined at the peptide sequence).

We investigated the molecular origin for the experimentally observed preferential swap. Proximity between free thiols has been suggested to be a key factor for disulfide bond swapping [59]. We tested if, under the application of force, the inter-sulfur distances in SD1 changed after the reduction of 1-4 or 2-8. We found that indeed, for the resulting four cases, inter-sulfur distances shifted considerably towards lower distance values upon force application of 500 pN (Fig.s 6A). The same general effect could not be observed for lower forces, presumably due to insufficient sampling. But still the attacking sulfur 2 maintained the tendency to occupy lower distance at a low force of 50 or 100 pN in an extra set of simulations started from a fully-elongated configuration. Consequently, force, by bringing the free thiol 2 into closer proximity to the attacked disulfide bond 1-4 appears to be an important factor defining one of the observed disulfide exchange reactions.

**Figure 6:**
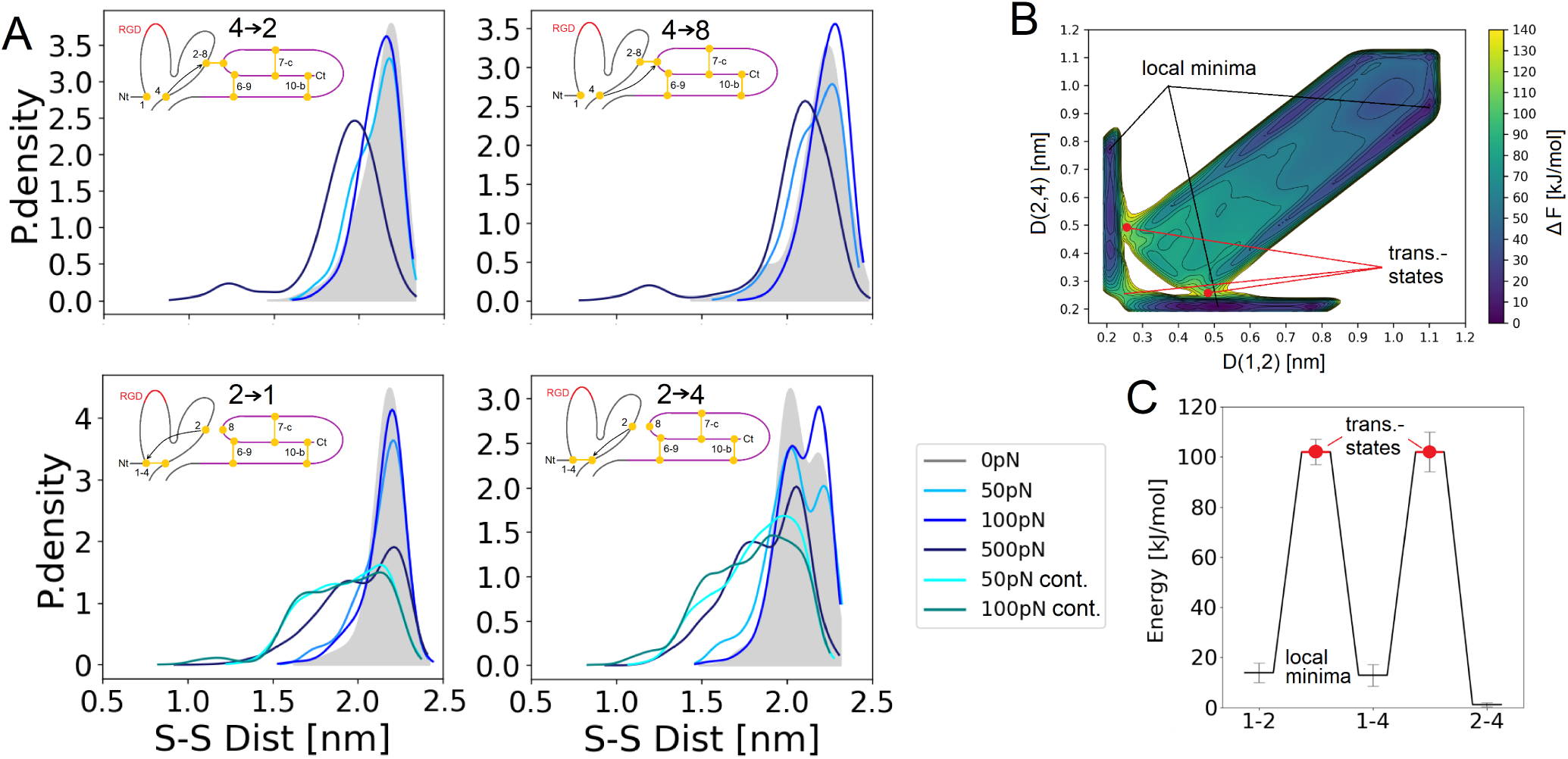
Force-mediated proximity along with thermodynamic affinity explains specific disulfide bond shuffling in vWF-C4. A: Inter-sulfur distance probability distributions when either the cysteine 4 or cysteine 2 is unpaired. Distance is presented for the indicated four sulfurs that could pair, under equilibrium (gray) and forced (blue shades) conditions. See all inter-cysteine distances in Figure S4. A schematic representation of the possible thiol disulfide exchange pathways is indicated at the inset. B: Free energy surface for the attack of the 2 free thiol to the 1-4 bond determined from QM/MM calculations (situation depicted at the bottom panels in A). X- and y-axis represent the inter-sulfur distance. Here, an external force of 166 pN was applied to the C4 termini. C: Energy minima and transitions barriers extracted from B. The black line marks the average result of all meta dynamics simulations. Error bars represent S.E.M. Transition states are indicated with the red dots. The energy minimum of 2-4 is significantly lower than those of 1-2 and 1-4, p < 0.02.

The proximity argument from MD simulations underlined the important role force plays bringing free thiols into close proximity. However, it did not resolve between the specific thiol being attacked. In particular, the free thiol 2 approached with similar probability both cysteines 1 and 4 while in mass spectrometry only binding to cysteine 4 could be verified (Fig 5). To compare the involved free energy changes of these swaps, we carried out QM/MM simulations using metadynamics and density-functional tight binding for the QM region. We concentrated on the attack of sulfur 2 to the 1-4 bond (Figure 6B), as this reaction can decide on the propensity of the RGD motif for binding to platelet integrins (see Fig. 6B–C). Figures 6B and C depict the two dimensional projection of the free energy surface and a simplified energy scheme, respectively, determined from a series of exchange reactions between sulfurs 1, 2 and 4.

While the transition barriers are equally high for both exchange options, the local minimum of configuration 2-4 lies significantly lower than those of all other permutations, a result that aligns very well with the experimental findings and that suggests 2-4 to be thermodynamically preferred over 1-2.

### Beyond C4: All vWF C domains are partially reduced

Motivated by the analysis of C4, we finally studied the prevalence of reduced disulfide bonds in the entire vWF-C domain family. Accordingly, the redox state of 17 disulfide bonds in the six C domains was quantified (Fig. 7). All 17 bonds were predominantly oxidised in ten healthy human donors (Fig. 7). A small fraction of these bonds, however, were on average 1-5 % reduced in the population of the vWF molecules. Interestingly, the observed frequency of bond reduction increases from C1 to C6 with greater proximity to the cysteine rich knot of the vWF. The 7-10 bond in C5 and C6 show the highest fraction of reduced vWF molecules, namely 6.5 and 7.3%, respectively. Consequently, not only C4 but all vWF-C domains have the potential to be labile.

**Figure 7:**
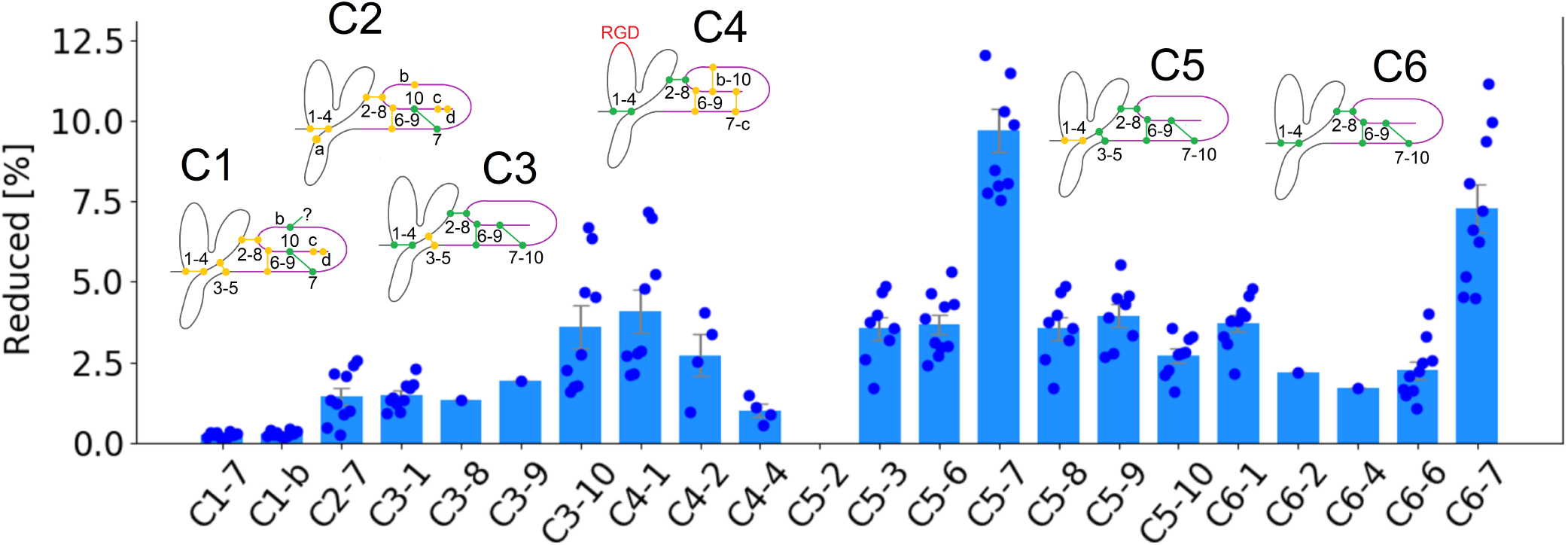
Redox states of 17 vWF C domain disulfides in ten healthy human donors. (5 male and 5 female, from 22–58 years old). The 17 bonds that have been analysed for redox state are colored in green. Blue dots are experimentally measured in individual donor blood. The bars and errors are mean ± S.E.M.

## Discussion

In this study, we investigated the structural and functional consequences of disulfide bond breakage and exchange for the vWF-C4 domain by molecular simulations, mass spectrometry and platelet binding assays, in equilibrium and under force. We started our analysis with the observation that a fully oxidized C4 domain is crosslinked by disulfides to an extent that locks the domain in a folded force-unresponsive state (Fig. 2). Partial reduction of specific bonds, however, enabled large conformational changes, thiol/disulfide exchange reactions, and mechano-redox regulation of integrin binding.

Two out of the five disulfide bonds present in C4, named 1-4 and 2-8, both located at the SD1 sub-region (Fig. 1) displayed a distinct behavior. Previous mass spectroscopy experiments [15] as well as our own experiments showed these two bonds to be reduced in blood (Fig. 3A). Their process of reduction itself is possibly catalyzed by a reducing agent such as glutathione or a redox enzyme such as protein-disulfide isomerase [9]. In addition, we find these two bonds to play a critical role for the structural integrity of the C4 domain, as their reduction leads to severe conformational rearrangements (Fig. 2). Hence, our combined findings provide evidence for the existence of at least two reduced redox states within the vWF-C4 domain and point towards a dramatic conformational alterations of this domain by the action of mechanical forces, such as those originating from the shear of the flowing blood, upon reduction.

The main biological function of the C4 domain is the reinforcement of platelet binding by interaction of its RGD motif with the platelet integrin receptor αIIbβ3. Platelet binding assays proved a decreased binding capacity in comparison to the fully oxidised C4 domain for those cases where either the 1-4 or 2-8 bond have been removed by mutation (Fig. 3B,C). Our MD simulations link this reduced binding ability to a flattening (1-4) or steric occlusion (2-8) of the RGD beta hairpin (Fig. 4). Thus, our data shows that reduction of these two specific disulfide bonds does not merely destabilize the C4 domain, but more importantly, compromises its main function of anchoring platelets to vWF in a force-dependent manner.

Blockage of free thiols has been reported to interfere with the binding of vWF to collagen [17] and to platelets [15]. Moreover, the methionine oxidation of vWF can regulate the protein’s force response and vice versa [60, 61]. Furthermore, auto-inhibition of the vWF for the initial binding of platelets is controlled by a disulfide bond in the vWF A2 domain [16]. Our study identifies the key molecular transitions in C4 that alter vWF-integrin binding. It thereby provides a structural and dynamic molecular explanation of how mechanical and redox stimuli jointly control vWF function. We propose that higher levels of oxidative stress decrease stable integrin-mediated platelet binding, while higher shear forces in blood push vWF into a less adhesive state at high oxidative stress conditions. In this scenario, processes with elevated oxidative stress levels, such as inflammation or immune response, could corroborate thrombotic disorders. Such mechano-redox crosstalk is emerging as a common theme also for other extracellular proteins such as integrins [18] or fibrinogen [19].

Disulfide bond reduction leads to the introduction of free thiols that could promote disulfide-bond exchange. In fact, our mass spectrometry findings demonstrate the formation of a new intra-C4 bond, between the cysteines 2 and 4 (Fig. 5), as well as across domains C3 and C4, between cysteine 10 of C3 and cysteine 1 of C4 (Fig. S2). Interestingly, disulfide bond swapping in the C2 domain is involved in vWF oligomerization [13]. It would be highly interesting to test if intermolecular disulfide swapping across C3 and C4 domains occurs between different vWF multimers. It would thereby promote vWF network formation and in this way complement non-covalent vWF assembly under shear conditions [62].

The mechanochemistry of disulfides has been intensively studied [63, 64]. Force-propelled intramolecular thiol/disulfide exchange has remained difficult to be observed in biological systems. Instead, disulfide bond swapping is considered to require enzymatic catalysis e.g. by PDI [9, 65]. To our knowledge, direct observation of a non-enzymati exchange has been so far only been possible for an immunoglobulin domain engineered for this purpose [63]. Here, we show that reduction of vWF in a stretched conformation brings specific thiols (thiol 2) close enough to one of the exchange candidates (1-4), such that swapping can occur (Fig. 6A). Proximity between the free sulfur and the attacked bond has indeed been suggested to be a key factor mediating disulfide-bond exchange [59, 66–68]. Also from an energetic point of view, the attack of the free thiol 2 on either bonded sulfur 1 or 4 is similarly likely, although a slight increased preference to 2-4 was observed (Fig.s 6B–C). The formation of the experimentally validated 2-4 bond would leave the RGD binding beta hairpin unprotected and prone to unfolding under the influence of external force, akin to the situation we observed when 1-4 was reduced (Fig. 4A). Thus, reduced integrin (and therefore platelet) binding of C4 due to complete flattening of the RGD binding site of vWF could also be a result of 2-8 to 2-4 disulfide bond exchange.

Our mass spectrometry data confirms the widespread occurrence of partially reduced disulfide bonds across all six C-domains (Fig. 7). Interestingly, the extent of reduction of disulfide bonds overall increases from the C1 to the C6 domain close to the C-terminal cystine-knot (CTCK). Homologous to C4, the C3, C5 and C6 domains exhibit partially reduced 1-4 bonds, suggesting pronounced force-induced unfolding of the N-terminal loop of these domains, with potential implications of stem formation [69].

In summary, we here establish the molecular determinants governing dynamic changes in the topology of vWF-C4’s disulfide bonds and the functional consequences of these changes on integrin-mediated platelet binding. Beyond structural integrity, mechano-sensitive disulfide-bond redox-control emerges as a prominent mechanism for the regulation of vWF function.

## Acknowledgement

We are grateful for financial support by the Klaus Tschira Foundation; the National Health and Medical Research Council of Australia (grant numbers 1110219, 1143400 and 1143398); the Senior Researcher Grant from the NSW Cardiovascular Research Capacity Program; the German Research Foundation (DFG) through the GRK 2450 grant FG, the state of Baden-Württemberg through bwHPC and the DFG through grant INST 35/1134-1 FUGG. We thank Mayukh Kansari for reparametrizing the sulfur-sulfur parameters for thiol-disulfide exchange with DFTB3.

## Notes

### Competing Interest Statement

The authors have declared no competing interest.

