## Supplementary material for "Dynamic Disulfide Bond Topologies in von-Willebrand-Factor’s C4-Domain Undermine Platelet Binding": Methods and Supplementary Figures

**Redox states of human vWF-C domain disulfides.** All procedures involving collection of human blood from healthy volunteers were conducted in accordance with the Human Research Ethics Committee of the University of Sydney (HREC 2014/244) and the Helsinki Declaration of 1983. Blood was collected by venesection using a 21 g butterfly Terumo needle from 10 healthy donors on no medications (5 male, 5 female, 18-57 years old). The first 5 mL of blood was discarded to avoid any thrombin that may have been generated around the needle insertion site and then drawn into tubes containing 3.2 % v/v sodium citrate. Plasma was prepared by twice centrifugation at 800 g for 20 min at room temperature. Dynabeads (2 mg, Life Technologies) were coated with 16 µg of polyclonal anti-vWF (Dako) antibodies in 1 mL of phosphate-buffered saline on a rotating wheel for 1 h at 22 °C and the excess antibody removed by washing 3 times with 1 mL of phosphate-buffered saline. Plasma (0.7 mL) was incubated with the 2 mg of coated beads on a rotating wheel for 1 h at 22 °C. The beads were collected using a magnet, excess plasma aspirated to reduce the volume, and incubated in 0.3 mL of 5 mM 2-iodo-N-phenylacetamide ($^{12}$C-IPA, Cambridge Isotopes) in phosphate-buffered saline containing 10% DMSO for 1 h at 22°C in the dark to alkylate unpaired CYS thiols in the proteins. The supernatants were aspirated, and the beads incubated with NuPAGE LDS sample buffer (Life Technologies) containing a further 5 mM $^{12}$C-IPA for 30 min at 60°C. The supernatants were resolved on SDS-PAGE. Purified recombinant vWF (19 µg) was incubated with 5 mM 12C-IPA in phosphate-buffered saline containing 10 % DMSO for 1 h at 22 °C in the dark and the protein resolved on SDS-PAGE.

The SDS-PAGE gels were stained with colloidal coomassie (Sigma) and the vWF or fibrinogen bands excised, destained, dried, incubated with 40 mM dithiothreitol and washed [1]. The gel slices were incubated with 5 mM carbon-13 isotope of IPA ($^{13}$C-IPA) for 1 h at 22 °C in the dark to alkylate the disulfide bonded cysteines, washed and dried before digestion of proteins with 12.5 ng/µl of trypsin (Promega) in 25 mM $NH_4CO_2$ overnight at 25 °C. Peptides were eluted from the slices with 5 % formic acid, 50 % acetonitrile (Fig. S3). Liquid chromatography, mass spectrometry and data analysis were performed as described [2, 3]. Cysteine labelled with $^{12}$C-IPA or $^{13}$C-IPA has a mass of 133.05276 or 139.07289, respectively. Cysteine-containing vWF peptides were analysed (Fig. S3 and Table S1). The different redox forms of the cysteine residues were quantified from the relative ion abundance of peptides labelled with $^{12}$C-IPA and/or $^{13}$C-IPA (Fig. S5). The data was searched for peptides containing free cysteine thiols and these were not detected, which indicates that alkylation of unpaired cysteine residues by $^{12}$C-IPA or $^{13}$C-IPA was complete in the proteins.

**Cloning and expression of human vWF.** Cysteine to Alanine substitutions at positions 2499 and 2533, and 2528 and 2570 were generated in full-length human vWF DNA using a Site-directed Ligase Independent Mutagenesis (SLIM) approach [4]. Briefly, the C2528A and C2570A mutants were created sequentially with C2528A being created first from a wild-type vWF template, followed by the fabrication of the C2570A mutation using the C2528A mutant as a template. Both mutations were confirmed by DNA sequencing. For expression in HEK293, constructs were cloned into pcDNA3.1 vector according to manufacturer's specifications and purified using PureLink Quick Plasmid Miniprep Kit (Machery-Nagel).

For transfection into HEK293, cells were grown at 37°C with 5 % $CO_2$ in Dulbecco's modified Eagle's medium, supplemented with 10 % fetal bovine serum (GE Healthcare) and penicillin-streptomycin antibiotics (Gibco). Cells at 60 % confluence were transfected using polyethylenimine at a ratio of 12.5 mg DNA: 50mg polyethylenimine (Polysciences) in OptiMEM media containing reduced serum (Life Technologies). Transiently transfected HEK293 cells were cultured at 37°C for 72 h before collection of conditioned media. Conditioned media (20 mL) was concentrated 10-fold using Snakeskin tubing (Thermofisher) and overlayed with polyethylene glycol 20,000 (Sigma). The concentrated proteins were dialysed overnight at 4°C into 20 mM Tris, 50 mM Na-Cl, pH 7.8 buffer and further concentrated using Ultracel spin filters with 100-kDa cut-off membrane (Millipore). Wild type vWF and the C4 mutants were resolved on 4-12 % Bis-Tris gels, transferred to a PVDF membrane and detected with polyclonal rabbit anti-human vWF-HRP (Dako). Quantification of the vWF proteins was performed using ELISA (Dako).

**Platelet adhesion studies.** Washed human platelets were perfused over vWF matrix using polydimethylsiloxane (PDMS) microfluidic channel devices. Briefly, microfluidic channels measuring 0.3 x 1.0 mm height x width were coated with polyclonal anti-vWF antibody (Dako, 40 µg/mL) and incubated in a humidified chamber at 4°C overnight. The following day, 20 µg/mL of recombinant wild-type or mutant vWFs (C2499/2533A or C2528/2570A) were added to vWF antibody pre-coated microfluidic channels and incubated at 37°C for 2 hours followed by blocking with 2 % BSA for 30 minutes.

Washed platelets were prepared from healthy donors according to established methods. Briefly, 10 mL of ACD anticoagulated whole blood was centrifuged at 200 g for 10 min without accelerator/brake. Platelet rich plasma (PRP) was collected and rested for 10 min before centrifugation at 1700 g for 5 min (brake level 7). Platelet pellet was resuspended in 4-5 mL of 1 x platelet wash buffer (1xPWB) containing 20 U/mL of Clexane and 0.01 U/mL of apyrase and allowed to rest for a further 10 min at 37°C. Platelets were pelleted by centrifugation at 1500 g for 5 min (brake at level 7) and resuspended in 1 ml of Tyrode's buffer. Platelets were rested for 30 min before use.

Calcein-FITC (1 mg/mL) labelled platelets (300-400 x 103 /mL) were perfused over the vWF-coated matrix at a shear rate of 1000 $s^{-1}$. Platelet adhesion was recorded using a Zeiss 880 confocal microscope with x40 water objective. Images were captured at 1 s/frame for 3 min. Platelets adhering to the matrix for at least 2s were considered adherent platelets. Rolling platelets were defined by their movement of >1 cell diameter in 10s following tethering to the matrix. Platelets moving <1 cell diameter in 10s were considered stationary platelets. The kinetics of platelet adhesion was analysed at 30s intervals. All images were quantitated using Image J analysis software [5].

**Mapping of disulfide bonds in human vWF-C domain.** Healthy donor plasma (0.7 mL) was incubated with polyclonal anti-vWF antibody coated Dynabeads on a rotating wheel for 1 h at 22°C. The beads were collected, excess plasma aspirated, incubated with NuPAGE LDS sample buffer for 30 min at 60°C and the supernatants resolved on SDS-PAGE. Gel slices containing the vWF were washed and dried before digestion with 12 ng/µL chymotrypsin (Roche Applied Science) in 25 mM $NH_4CO_2$ containing 10 mM $CaCl_2$ for 16 h at 25°C with vortexing. The vWF was further digested with 12 ng/µL trypsin (Promega) for 4 h at 37°C with vortexing. Reactions were stopped by adding 5 % (v/v) formic acid and peptides eluted from the gel slices with 5 % formic acid and 50 % (v/v) acetonitrile. Peptides in 0.1 % formic acid (final volume 12 µL) were resolved on a 35 cm × 75 µm C18 reverse phase analytical column using a 2–35 % acetonitrile gradient over 22 min at a flow rate of 300 nL/min (Thermo Fisher Scientific Ultimate 3000 HPLC). Peptides were ionized by electrospray ionization at +2.0 kV and analysed on a Q-Exactive Plus mass spectrometer (Thermo Fisher Scientific) using collision-induced dissociation fragmentation. Disulfide-linked peptides were searched against human vWF sequence using Byonic analysis software (Version 3.9-7, Protein Metrics) with a false discovery rate of 1 %. Disulfide-linked peptides were manually inspected for accuracy. Precursor mass tolerance and fragment tolerance were set at 10 ppm and 0.6 Da, respectively. Variable modifications were defined as oxidized Met, oxidized cysteine (mono, di and tri) and glutathionylated cysteine with full trypsin and chymotrypsin cleavage of up to three missed cleavages.

**Equilibrium MD simulations.** Different conformations of the C4 domain (fragment 2493-2577 of the vWF sequence, i.e. 85 residues) were taken from previous MD simulations [6] in a randomised fashion. We removed individual inter-cysteine disulfide bonds, resulting in 6 different cases: one with the protein fully oxidised (fo) and five more with one of the disulfide bonds removed (1-4, 2-8, 6-9, 7-c, b-10) (Fig. 1 and 2 ofthe main text). At least ten different starting conformations were considered for the fo as well as for any of the reduced cases. The C4 domain was put into solution with ∼100.000 explicit water molecules, enough to accommodate C4 in a potentially extended conformation. 0.15 mol/L Na-Cl ions were added to the system with excess ions assuring a net neutral state of charges. We performed an energy minimisation and subsequently 1 ns of solvent-equilibration with position restrains applied on the heavy atoms of the protein with an elastic constant of 1000 kJ/mol/nm in each spatial dimension. Position restraints were removed and, for each structure, production runs of 140–210 ns were considered. The first 20 ns of every run were discarded to further equilibrate the system, resulting in at least 1.4 µs of total simulation time for each system.

**Force-probe MD simulations.** N- and C-terminus of C4 were pulled away from each other applying a constant external force to both termini in opposite directions. This procedure was performed for three different force regimes, first at ∼500 pN (300 kJ/mol/nm), thereafter at ∼100 pN (60 kJ/mol/nm) and ∼50pN (30 kJ/mol/nm). At least 10 independent simulation runs were considered for each of the six cases. Starting structures of the protein were taken at random times from the equilibrium simulations. Subsequently, the protein was solvated by 55.000–135.000 water molecules in a simulation box of dimensions $60\times8.2\times8.2\,nm^3$ for the 500 pN 1-4 case, $50\times6\times6\,nm^3$ for 50 pN and 100 pN 1-4, $25\times8.2\times8.2\,nm^3$ for all other 500 pN cases and $30\times6\times6\,nm^3$ for all other 50 pN and 100 pN cases (long enough in the x-axis to accommodate the unfolded domain). Subsequently this system was also neutralised by excess ions and 0.15 mol/L of Na-Cl was added. The same protocol and parameters were used as in the equilibrium simulations. For each system, at least 1 µs of cumulative simulation time was generated with 70–200 ns run time for each simulation. In addition, we extracted fully extended molecular configurations from the 500 pN 2-8 case at random times and used them as starting structuress for 10 new simulations of at least 100 ns length applying a force of 50 pN and 100 pN in a $50\times6\times6\,nm^3$ box. We employed the same energy minimisation and solvation procedure as we did for the configurations extracted from equilibrium.

**MD simulation details.** MD simulations were carried out by the GROMACS software suite versions 2018 and 2020 [7], making use of the LINCS algorithm to constrain protein bonds involving hydrogens [8] and of the SETTLE algorithm [9] to constrain both angular and bond vibrations of water, enabling a 2 fs time step to propagate the Leap-Frog algorithm Periodic boundary conditions were imposed. Neighbors were treated with the Verlet cutoff-scheme [10]. Electrostatics were computed using the Particle Mesh Ewald method [11, 12]. Short-range interactions were modelled with a Lennard-Jones potential, employing a cutoff radius of 1.1 nm and switching the force to zero from 1 to 1.1 nm. Temperature and pressure were maintained constant at 300 K and 1 bar, respectively, by employing the Berendsen thermo- and barostat [13] during the solvent-equilibration ($\tau_T = 0.5\,ps$, $\tau_P = 5\,ps$), and the V-rescale thermostat [14] as well as the Parinello-Rahman barostat [15] during the production runs ($\tau_T = 2.5\,ps$ and $\tau_P = 5\,ps$). Protein and non-protein atoms were coupled separately to the thermostat. Force field parameters were taken from the amber99sb-ildnp-star suite for the protein [16, 17]. The TIP3P water model [18] and default amber ion parameters were used.

For subsequent analysis, coordinates $(x, y, z)$ of every atom were extracted from the trajectories after removing unwanted molecular jumps over the boundaries of the simulation box.

**The beta hairpin angle.** C-α atom positions were selected for CYS2499 (1 of the 1-4 bond), GLY2508 (from the RGD motif), and CYS2533 (4 of the 1-4 bond) in every frame. The resulting angle formed by these three atoms was calculated with an in-house python script. This angle was considered as a measure for the degree of opening of the beta hairpin containing the RGD motif.

**The Ramachandran analysis.** Positions of the N, C-α and C backbone atoms of the RGD motif (residues 2507, 2508 and 2509) as well as the C atom of adjacent residues 2506 and the N atom of residue 2510 were extracted from the simulations. The dihedral angles φ(involving C,N,C-α ,C) and ψ(corresponding to N,C-α ,C,N) were accordingly calculated with an in-house python script.

**Binding of C4 to platelet integrin.** We used the AlphaFold2 software [19, 20] implemented in ChimeraX [21] to determine a likely binding configuration of vWF-C4 bound to $\alpha$IIb$\beta$3 integrin. Thereafter, we superposed the C4 configurations from the simulations with that predicted by AlphaFold2 via least-square-fitting of the backbone atom positions of the RGD carrying beta hairpin (residues 2500 to 2514). After receiving atomic positions from alphafold, the fit could be performed using the GROMACS-trjconv algorithm suite.

**Atomic clashes.** We monitored the number of atomic clashes (distances below 0.3 nm) between heavy atoms of the superposed conformation of the C4 domain and the backbone atoms of the integrin predicted by AlphaFold2. We associated higher number of clashes with poorer quality of fit. To calculate the contacts, we employed an in-house python script. We computed the number of clashes $N$ separately for the SD1 and SD2 subdomains. In the main text we present $N$(SD1), $N$(SD2), and $N$(SD1)/[1 + $N$(SD2)].

**Sulfur Proximity.** Sulfur atomic positions were extracted from the trajectories using the GROMACS-traj tool. Inter-atomic distances could then directly be determined via an in-house python script.

**P-values.** From the MD time-series, we took $n$ auto-correlated data points (tipically spaced by 0.4 ns). Distributions, means, standard deviations and standard errors on the mean were presented in the main text. Where indicated, P-values were computed with t-tests on the basis of the effective number of uncorrelated frames calculated by the formula described in [22]:

$$n_{\text{eff}} = \frac{n}{1 + 2\sum_{k=1}^{n_c}\left(1 - \frac{k}{n}\right) r_k}, \qquad (1)$$

with $r_k$ the auto-correlation function estimated from the data and $n_c$ the number of time frames until first zero transition of $r_k$. If several independent simulations were investigated, describing the same system, the median of all $n_{eff}$ was taken as the best estimate for all simulations.

**QM/MM simulations.** **Equilibration:** Seven structures with high mutual RMSD values were selected from the Force-probe MD runs to serve as initial structures for QM/MM metadynamics simulations. Each structure was equilibrated in an NVT ensemble at 300 K with the Bussi thermostat [14] for over 500 ps using the leap-frog integrator with a time step of 1 fs. A local version of Gromacs 2020 patched with Plumed 2.5.1 and interfaced with DFTB+ 19.1 was used for the simulations [7, 23, 24]. The side chains of CYS 1, CYS 2 and CYS 4 up to $C_\beta$ were included in the QM region, to be described with the semi-empirical density functional theory method DFTB3 using the 3OB parameter set [25, 26]. It has been shown, that DFTB3 using the 3OB parameter set [26] does not describe thiol-disulfide exchange reactions with sufficient accuracy [27]. Thus, we performed a specific reaction parameterization (SRP) for the sulfur-sulfur parameters in Ref. 28. The SRP which was fitted to G3B3 data was used in this work. Each covalent bond connecting a QM with an MM atoms was treated with a link atom, i.e. a hydrogen atom was inserted along the $C_\alpha$–$C_\beta$ bond; the complete QM region then contained 15 atoms. The rest of the system was described with the Amber99SB-ILDN force field [16] and the TIP3P water model [18]. During the simulation, a constant pulling force of 500 kJ/mol/nm (830 pN) along the x-axis in the opposite directions was applied to the centers of masses of the terminal amino acids. The electrostatic interactions between the QM and MM regions were scaled down by a factor of 0.75 (inverse square root of the optical dielectric constant) to compensate for the missing electronic polarization of the MM environment as suggested by Leontyev & Stuchebrukhov [29, 30].

**Metadynamics:** In the next step, well-tempered multiple walker metadynamics was used to obtain the potentials of the mean force of the thiol/disulfide exchange reactions [31–33]. The pulling force was reduced to 100 kJ/mol/nm (166 pN). A separate metadynamics simulation was performed using each of the different starting structure (following the equilibration) with the force of 166 pN. Each simulation involved 16 walkers simulated for at least 18.75 ns at 300 K with the Bussi thermostat yielding a total simulation time in excess of 300 ns per simulation, and more than 2 µs of QM/MM metadynamics in total. The three distances between the sulfur atoms were used as reaction coordinates, and gaussian bias potentials were added every 500 fs; the potentials of all walkers were exchanged every 1000 fs. The initial gaussian height was set to 1.0 kJ/mol, the width to 0.2 Å and the bias factor to 50.

**Additional restraints:** Harmonic restraints with a force constant of 100.000 kJ/mol/nm were applied to the distances S2–S1 and S2–S4 above 1.1 Å and to S1–S4 above 6 Å, to reduce the volume of configurational sample space. To avoid chemically wrong reactions due to accumulated biasing potentials on the S–S distances, the same restraints described in Ref. 34 and 28 were used. These include restraints on the sum of bond switching functions of the three S–S distances, on the sum of the S–S distances, on the individual S–S distance coordination numbers, on the mean coordination numbers of each carbon with its two protons, and on each sulfur–hydrogen distance.

### Supplementary Figures

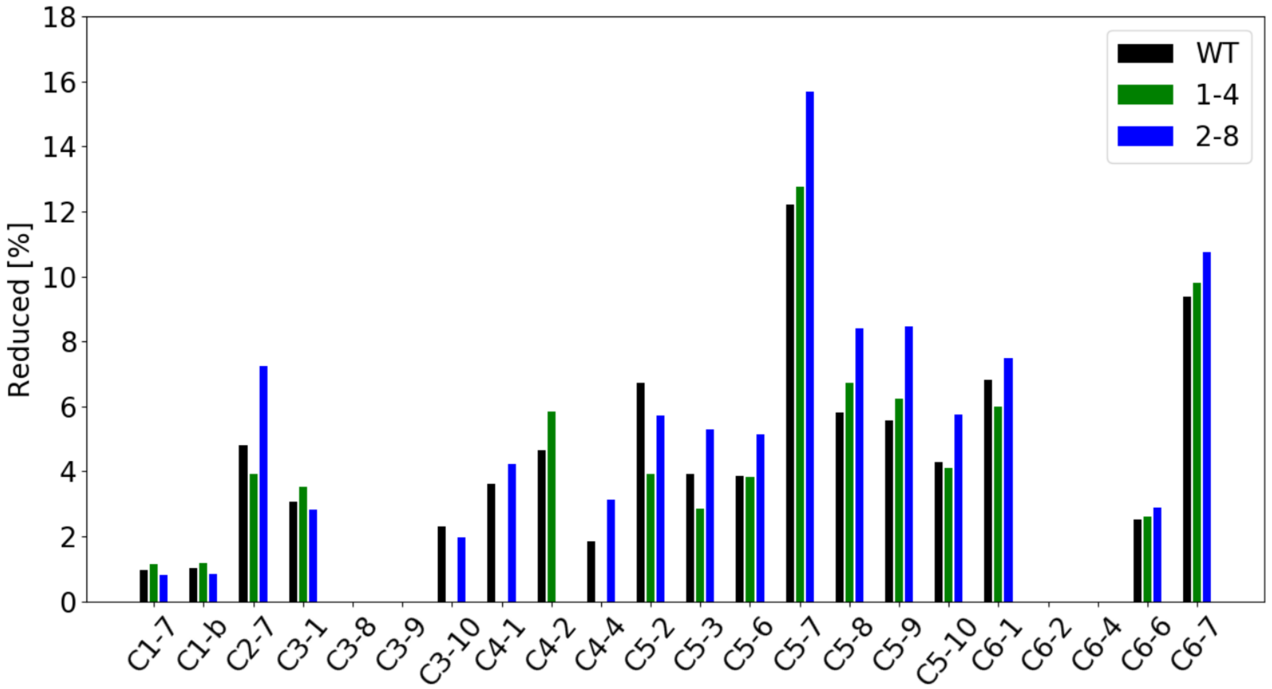


**Figure 1:** ***Redox states of C domain disulfides in wild-type (WT) and C2499,2533A (1-4) and C2528,2570A (2-8) vWF mutants:*** *Elimination of the C4 1-4 bond does not appreciably change the redox state of the C4 2-8 bond, and vice versa. The same holds true for all other disulfide reduction percentages in the C domain region of vWF.*

| S-S bond | CYS | Domain | Peptide |
|---|---|---|---|
| 2304-2325 | 2304 | C1: **7**-10 | APT**C**GLCEVAR |
| 2307-? | 2307 | C1: **b**-? | APTCGL**C**EVAR |
| 2375-2394 | 2375 | C2: **7**-10 | VSPPS**C**PPHR |
| 2431-2453 | 2431 | C3: **1**-4 | V**C**VHR |
| 2448-2490 | 2490 | C3: 2-**8** | SGFTYVLHEGE**C**CGR |
| 2473-2491 | 2491 | C3: 6-**9** | SGFTYVLHEGEC**C**GR |
| 2477-2494 | 2494 | C3: 7-**10** | **C**LPSACEVVTGSPR |
| 2499-2533 | 2499 | C4: **1**-4 | CLPSA**C**EVVTGSPR |
| 2528-2570 | 2528 | C4: **2**-8 | SVGSQWASPENP**C**LINECVR |
| 2499-2533 | 2533 | C4: 1-**4** | SVGSQWASPENPCLINE**C**VR |
| 2600-2640 | 2600 | C5: **2**-8 | TVMIDV**C**TTCR |
| 2603-2619 | 2603 | C5: **3**-5 | TVMIDVCTT**C**R |
| 2582-2605 | 2605 | C5: 1-**4** | **C**MVQVGVISGFK |
| 2624-2641 | 2624 | C5: **6**-9 | TT**C**NPCPLGYK |
| 2627-2644 | 2627 | C5: **7**-10 | TTCNP**C**PLGYK |
| 2600-2640 | 2640 | C5: 2-**8** | EENNTGE**C**CGR |
| 2624-2641 | 2641 | C5: 6-**9** | EENNTGEC**C**GR |
| 2627-2644 | 2644 | C5: 7-**10** | **C**LPTACTIQLR |
| 2649-2676 | 2649 | C6: **1**-4 | CLPTA**C**TIQLR |
| 2671-2715 | 2671 | C6: **2**-8 | DETLQDG**C**DTHFCK |
| 2649-2676 | 2676 | C6: 1-**4** | DETLQDGCDTHF**C**K |
| 2693-2716 | 2693 | C6: **6**-9 | VTG**C**PPFDEHK |
| 2701-2719 | 2701 | C6: **7**-10 | **C**LAEGGK |

**Table 1:** *VWF cysteine containing peptides analysed by HPLC and mass spectrometry. Cysteine numbering is according to UniProt identifier P04275 for human von Willebrand factor. The cysteine residues of the disulfide that were measured are underlined in the peptide.*

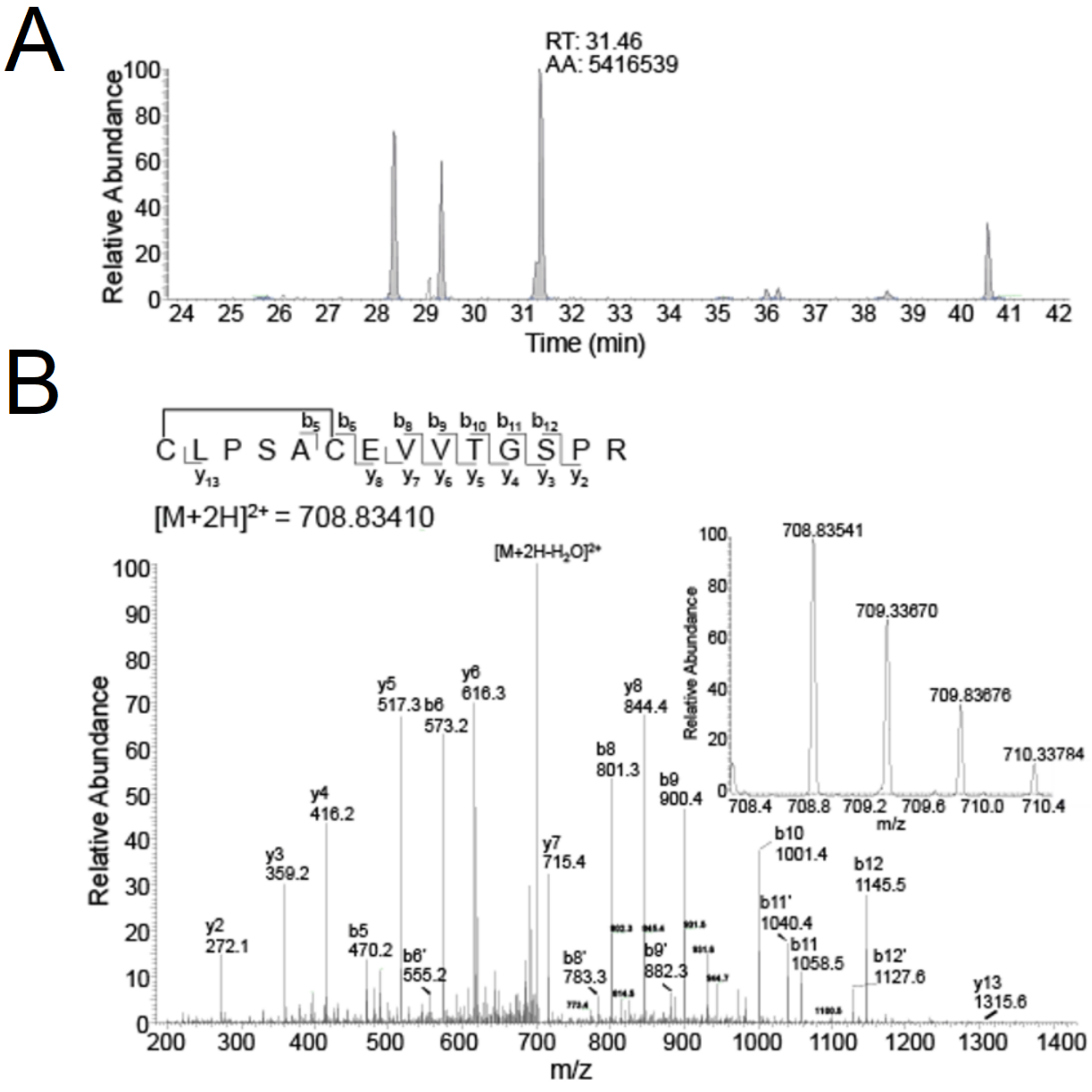


**Figure 2:** ***Identification of the vWF C3/C4 domain C2494-C2499 disulfide bond:*** *(a) HPLC resolution of the C3/C4 domain CLPSACEVVTGSPR peptide containing the C2494-C2499 disulfide bond in vWF from healthy donor plasma. (b) Representative tandem mass spectra of the CLPSACEVVTGSPR peptide. The accurate mass spectrum of the peptide is shown in the inset (observed [M+2H]$^{2+}$ = 708.83541 m/z and expected [M+2H]$^{2+}$ = 708.83410 m/z) .*

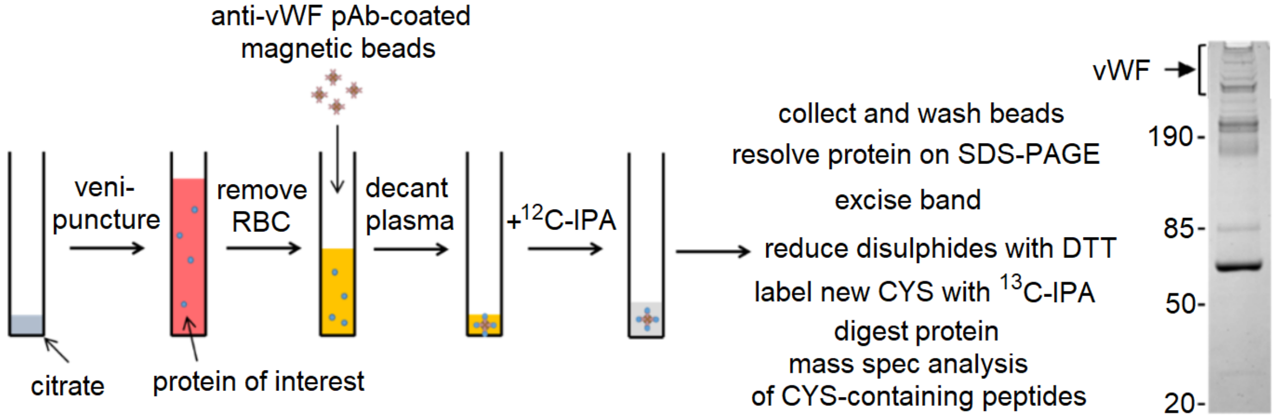


**Figure 3:** ***Disulfide bond determination:*** *Blood from healthy donors was drawn by venipuncture into citrate as anti-coagulant, plasma prepared by centrifugation and vWF collected on polyclonal anti-vWF antibody-coated magnetic beads. The vWF was digested with trypsin and chymotrypsin and peptides quantified by HPLC and identity established by mass spectrometry.*

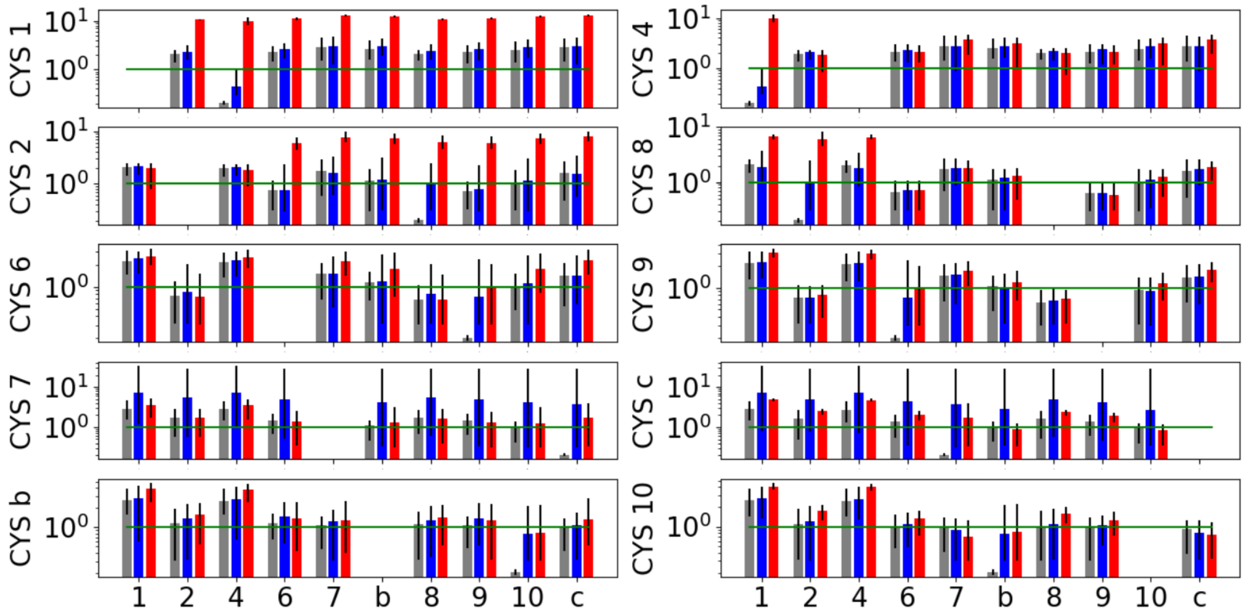


**Figure 4:** ***Inter-sulfur distance for different redox states of C4:*** *Shown are the average distances between sulfur x and y in the C4 domain under fully oxidised equilibrium conditions (grey bars), reduced equilibrium conditions where the y-axis sulfur also indicates which disulfide bridge underwent reduction (blue bars) as well as the distances for the reduced states under 500 pN force (red bars). Black error bars indicate the registered maximum and minimum distance. Green horizontal lines mark the 1 nm distance.*

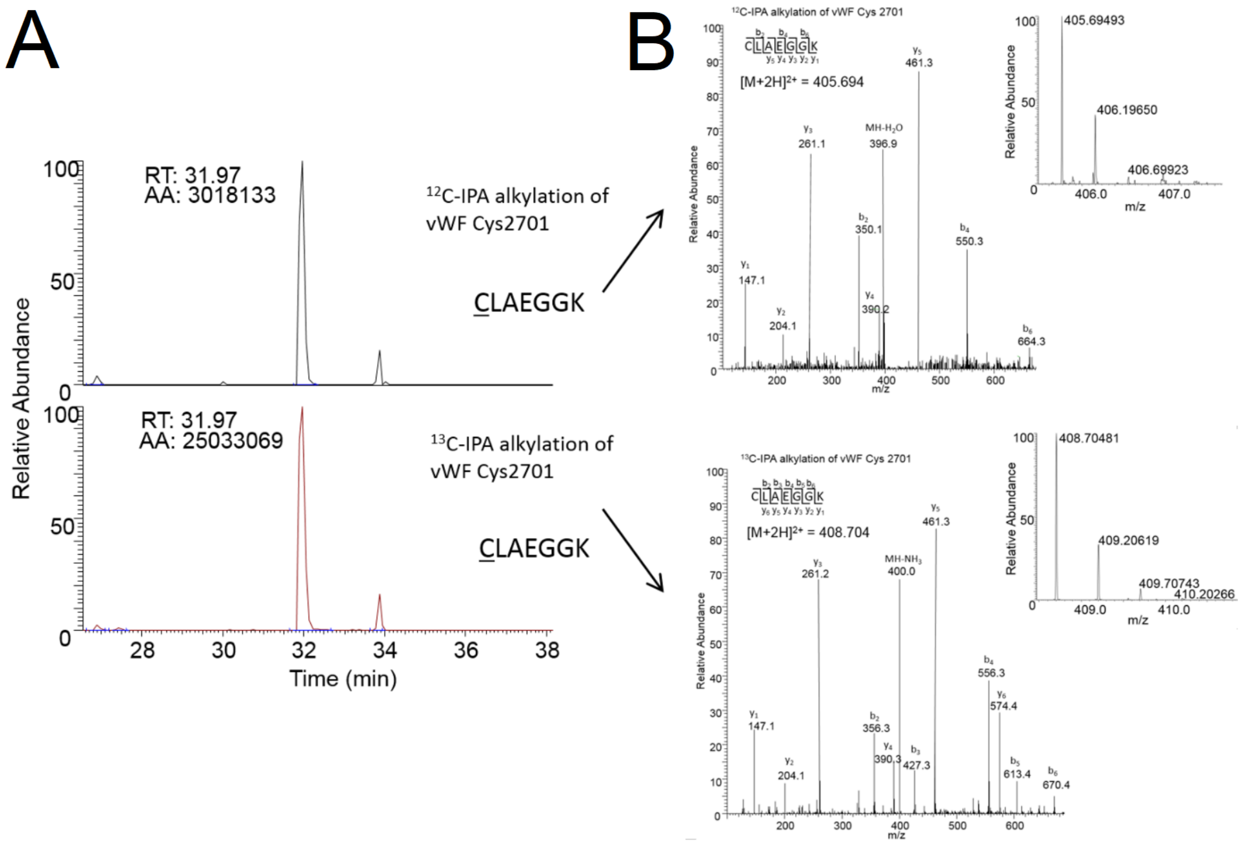


**Figure 5:** ***Differential cysteine alkylation of the vWF C6 domain CYS2701 residue and peptide analysis:*** *(a) HPLC resolution of the C6 domain CLAEGGK peptide containing CYS2701 labelled with either 12C-IPA (upper trace) or 13C-IPA (lower trace). (b) Representative tandem mass spectra of the C6 domain CLAEGGK peptide. The upper and lower traces are examples of 12C-IPA or 13C-IPA alkylation of CYS2701, respectively. The accurate mass spectrum of the peptide is shown in the insets (upper trace, observed* $[M+2H]^{2+} = 405.69493$ *m/z and expected* $[M+2H]^{2+} = 405.694$ *m/z; lower trace, observed* $[M+2H]^{2+} = 408.70481$ *m/z and expected* $[M+2H]^{2+} = 408.704$*) m/z.*
